# Transposable elements contribute to the establishment of the glycine shuttle in Brassicaceae species

**DOI:** 10.1101/2022.12.06.519256

**Authors:** Sebastian Triesch, Alisandra K. Denton, Jacques W. Bouvier, Jan P. Buchmann, Vanessa Reichel-Deland, Ricardo Nuno Ferreira Martins Guerreiro, Noah Busch, Urte Schlüter, Benjamin Stich, Steven Kelly, Andreas P.M. Weber

## Abstract

C_3_-C_4_ intermediate photosynthesis has evolved at least five times convergently in the Brassicaceae, despite this family lacking *bona fide* C_4_ species. The establishment of this carbon concentrating mechanism is known to require a complex suite of ultrastructural modifications as well as changes in spatial expression patterns, which are both thought to be underpinned by a reconfiguration of existing gene-regulatory networks. However, to date, the mechanisms which underpin the reconfiguration of these gene networks are largely unknown. In this study, we used a pan-genomic association approach to identify genomic features that could confer differential gene expression toward the C3-C4 intermediate state by analysing eight C3 species and seven C3-C4 species from five independent origins in the Brassicaceae. We found a strong correlation between transposable element (TE) insertions in *cis*-regulatory regions and the C_3_-C_4_ intermediacy. Specifically, our study revealed 113 gene models in which presence of a TE within a gene correlates with C_3_-C_4_ intermediate photosynthesis. In this set, genes involved in the photorespiratory glycine shuttle are enriched, including the glycine decarboxylase P-protein whose expression domain undergoes a spatial shift during the transition to C_3_-C_4_ photosynthesis. When further interrogating this gene, we discovered independent TE insertions in its upstream region which we conclude to be responsible for causing the spatial shift in *GLDP1* gene expression. Our findings hint at a pivotal role of TEs in the evolution of C_3_-C_4_ intermediacy, especially in mediating differential spatial gene expression.

## Introduction

C_4_ photosynthesis has convergently evolved more than 60 times in flowering land plants (Sage et al. 2012). C_4_ photosynthesis functions as a biochemical carbon concentrating mechanism that reduces the rate of photorespiration and thereby increases photosynthetic efficiency. Species that perform C_4_ photosynthesis are mainly found in warm, dry and high-light environments in which leaf internal CO_2_ are frequently low and by extension, the oxygenation to carboxylation ratio of Rubisco is elevated (Betti et al. 2016, Sage et al. 2012). Although C_4_ photosynthesis has evolved independently in multiple disparate plant lineages, the complexity of the required anatomical, biochemical, and developmental adaptations make engineering C_4_ photosynthesis a difficult undertaking.

Plants that exhibit C_3_-C_4_ intermediate phenotypes are promising research subjects to study the early steps towards C_4_ photosynthesis (Bellasio & Farquhar 2019, Kennedy & Laetsch 1974, Lundgren 2020, Schlüter & Weber 2016). C_3_-C_4_ intermediate species exhibit specialized anatomical traits and they differ from C_4_ species as they do not possess a fully integrated C_4_ cycle. C_3_-C_4_ intermediate traits are characterized by a lowered CO_2_ compensation point (CCP), chloroplast and mitochondria-rich bundle-sheath cells (BSC) and, in some cases, an increased vein density (Christin et al. 2011, Dengler et al. 1994, Schlüter et al. 2017). A further trait that is commonly shared between C_3_-C_4_ intermediate species from independent origins is the photorespiratory glycine shuttle, sometimes referred to as C_2_ photosynthesis (reviewed in Schlüter & Weber (2016)). This shuttle relies on the BSC-specific decarboxylation of photorespiratory glycine, leading to an elevated CO_2_ concentration around Rubisco. By extension, this increased partial pressure of CO_2_ around the site of its fixation leads to a higher frequency of the Rubisco carboxylation reaction compared to oxygenation reactions, thereby suppressing photorespiration and resulting in decreased CCP (Kennedy & Laetsch 1974, Monson & Edwards 1984, Schlüter et al. 2017).

Changes in the spatial and temporal patterns of gene expression are crucial for the evolution of C3-C4 intermediate photosynthesis (Hibberd & Covshoff 2010, Reeves et al. 2017). Previously, it has been shown that the BSC-specific decarboxylation of glycine is caused by the differential localization of the glycine decarboxylase complex (GDC). In C_3_-C_4_ intermediate species from the genera *Moricandia, Flaveria* and *Panicum*, the P-protein of the GDC is only observed in BSC mitochondria, but not in mesophyll cell (MC) mitochondria (reviewed in Schulze et al. (2016)). This is a notable example of convergent evolution, as these species belong to the distant families Brassicaceae, Asteraceae and Poaceae, respectively. In these plants, loss of the GDC P-protein from the MC restricts glycine decarboxylation to the BSC in C_3_-C_4_ intermediate species (Morgan et al. 1993, Rawsthorne et al. 1988, Schulze et al. 2016). However, the exact mechanism by which this is achieved differs in different species. For instance, in C_3_ *Flaveria*, the gene encoding the GDC P-protein (GLDP) is present in two differentially regulated copies, *GLDPA* and *GLDPB*. In C_3_-C_4_ intermediate *Flaveria* species, the ubiquitously expressed *GLDPB* is downregulated compared to C_3_ *Flaveria* species, whereas the BSC-specific *GLDPA* is highly expressed (Schulze et al. 2013). In contrast, in C_3_-C_4_ intermediate *Moricandia*, the differential expression of *GLDP* is thought to be mediated by the loss of one gene copy and a change in regulation of the other copy. Specifically, in C_3_-C_4_ intermediate Brassiceae species, *GLDP2* is absent and *GLDP1* was reported to be differentially expressed by loss of a potential *cis*-element called M-Box. The M-Box element in the *Arabidopsis thaliana GLDP1* promoter confers a low-level expression in both MC and BSC and is absent from the upstream region of *GLDP1* in C_3_-C_4_ intermediate *Moricandia* species. A second *cis*-element, the V-Box, was shown to confer high levels of expression in the BSC and is present in all analyzed Brassicaceae *GLDP1* promoter sequences to date (Adwy et al. 2015, 2019). Thus, there are multiple mechanisms through which *GLDP1* expression can be changed from being ubiquitously expressed in the leaf, to being BSC-specific in C_3_-C_4_ plants.

Structural variation can originate due to the activity of mobile genetic elements. In plants, transposable elements (TEs) comprise a large fraction of mobile genetic elements and contribute substantially to genome size variation (Lee & Kim 2014) and have substantial effects on the expression of genes (Hirsch & Springer 2017). TEs can be divided into two classes (Wicker et al. 2007) based on their transposition mechanisms: Class I transposons proliferate via a “copy-and-paste “ mechanism involving an RNA intermediate, whereas Class II transposons transpose directly via a “cut-and-paste “ mechanism. Due to their impact on structural variation, it has been frequently proposed that TEs can play a part in genome evolution and the evolution of novel genetic and phenotypic features (Buchmann et al. 2012, Feschotte 2008, Qiu & Köhler 2020, Wicker et al. 2007). Decades ago, Britten & Davidson (1971) put forward the idea that co-option of mobile sequences containing gene regulatory elements can connect genes to the same gene regulatory networks. The co-option of TEs for regulatory purposes is called “exaptation “ (Brosius & Gould 1992). In the present day with the vast amount of genomic data available, a deeper understanding of the role of transposable elements in genetic regulation allows linking genomic mechanisms with the evolution of complex traits.

TEs can rewire gene regulatory networks using different modes of action and influence the interplay of regulatory proteins (*trans*-elements) and the DNA sequences they are binding to (*cis*-elements). One such mode of action is the exaptation of a *cis*-regulatory element (CRE) from a separate gene (Fig 1). If the CRE inside a TE is copied from one gene and retained by the other gene, both genes become controlled by a mutual CRE and are thus connected by a shared gene regulatory network (Fig 1, B). In contrast to this scenario, it is also possible that TE integration into a CRE can suppress its function, either by interrupting the CRE sequence or altering the chromatin state of the respective CRE locus (Fig. 1, C) (Feschotte 2008). A further possibility is the *de novo* generation of new CRE by point mutations in TEs (Fig. 1, D). New CREs, e.g. a 10-mer promoter element, can arise by random point mutations in between 700,000-4.8 million years (Behrens & Vingron 2010).

**Figure 1.**
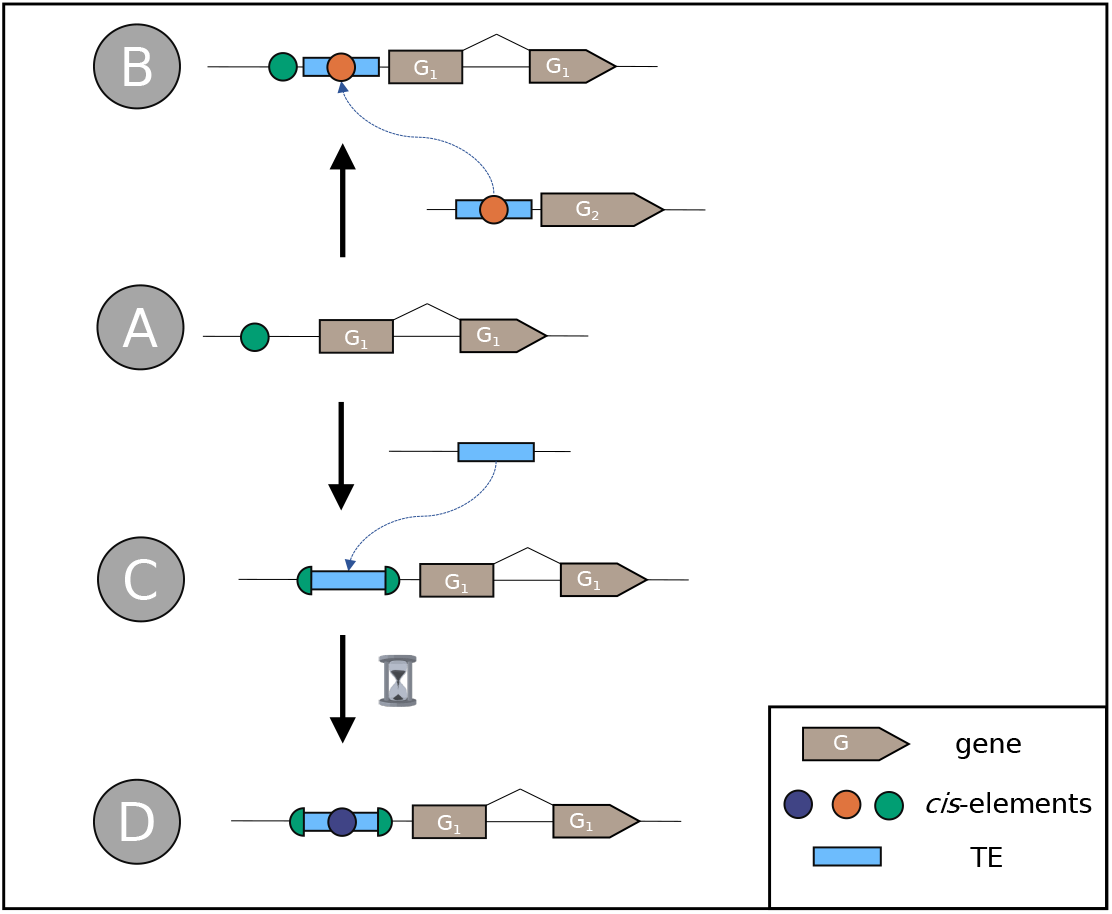
Schematic illustration of gene regulation rewiring by transposable element (TE) exaptation. A: The hypothetical gene G1 is controlled by a *cis*-regulatory element (CRE; green dot). B: Gene G2 is regulated by a different CRE (orange dot) located within a TE (blue box). Upon transposition of the TE to the upstream region of G1, G1 might co-opt the function of the orange CRE, thus connecting G1 and G2 to the same gene regulatory network. C: TE transposition can also lead to destruction or suppression of the CRE. D: During TE decay, new CREs (blue dot) might occur through accumulation of point mutations.

Several examples for the role of TEs in rewiring gene regulatory networks in plants have been reported. In rice, the *mPing* DNA transposon was found preferentially in the 5 ‘ region and was associated with the upregulation of stress response genes (Naito et al. 2009). In Brassicaceae, the evolution of heat-tolerance was linked to the activity of *Copia* retrotransposons containing heat-shock factor binding elements (Pietzenuk et al. 2016). Furthermore, TEs were also found to be associated with endosperm development, e.g. the distribution of the PHEREs1 MADS-box transcription factor binding motifs by *Helitron* transposons in *Arabidopsis thaliana* (Batista et al. *2019). The Youren* miniature inverted-repeat TE (*MITE*) was shown to be transcribed in rice endosperm, putatively mediated by a NUCLEAR FACTOR Y binding motif in the vicinity of the 5 ‘ terminal inverted repeat (TIR) of *Youren* (Nagata et al. *2022)*.

Previously, it has been shown that TEs play a significant role in the evolution of C_4_ photosynthesis in maize. For instance, by analyzing 40 C_4_ gene orthologs between rice and maize for the presence of BSC-specific promoter motifs, Cao et al. (2016) identified over 1,000 promoter motifs that were differentially distributed between C_3_ and C_4_ orthologs, of which more than 60 % were found to be associated with TEs and potentially co-opted by TE integration. These motifs may originate from non-photosynthetic genes and transposed to C_4_ genes, which connected gene regulatory networks. The authors showed that TEs play a significant role in the evolution of C_4_ photosynthesis in maize. However, the study of Cao et al. (2016) focused on evolutionary distant grasses, which makes it difficult to draw conclusions about the early evolutionary events towards C_4_ photosynthesis.

In the present study, we test whether TE insertions are involved in decisive steps of the evolutionary establishment of C_3_-C_4_ intermediate photosynthesis. To do this, we focused on the Brassicaceae family which exhibits at least five independent origins of C_3_-C_4_ intermediate photosynthesis (Guerreiro et al. 2023, *Schlüter et al. 2022) and contains multiple important and well-studied model plant species such as Arabidopsis thaliana, Arabis alpina* as well as relevant crop and vegetable plants such as *Brassica oleracea* and *Diplotaxis tenuifolia* (arugula).

We performed a pan-genomic association study to analyze the TE landscape of 15 Brassicaceae species. We tested for correlations between TE positions and the presence of C_3_-C_4_ intermediate traits. Specifically, we tested for correlations between the presence or absence of upstream co-occurring TEs with the CO_2_ compensation point. In this unbiased approach, we aimed at finding genes that retained upstream TEs selectively only in C_3_-C_4_ intermediate plants. Based on the results of this analysis, we examined the upstream regions of relevant photorespiratory genes in closer detail to assess the potential role that TE insertions have played during establishment of C_3_-C_4_ photosynthesis traits. In doing so, we present evidence that the insertion of TEs in *cis*-regulatory regions of key genes is associated with the evolution of C_3_-C_4_ photosynthesis in the Brassicaceae.

## Materials and Methods

### Genomes and carbon compensation points

The genomes of *Brassica gravinae* (Bg), *Brassica tournefortii* (Bt), *Carrichtera annua* (Ca), *Diplotaxis erucoides* (De), *Diplotaxis tenuifolia* (Dt), *Diplotaxis viminea* (Dv), *Hirschfeldia incana* (accessions HIR1 and HIR3), *Moricandia nitens* (Mn) and *Moricandia suffruticosa* (Ms) were obtained from Guerreiro et al. (2023). The genome of *Arabis alpina* (Aa) was obtained from Jiao et al. (2017). The genome of *Arabidopsis thaliana* (At) was obtained from Lamesch et al. (2012). The genome of *Moricandia arvensis* (Ma) and *Moricandia moricandioides* (Mm) were obtained from Lin et al. (2021). The genome assembly for *Brassica oleracea* (Bo) was obtained from Parkin et al. (2014). The genome for *Gynandropsis gynandra* (Gg) was obtained from Hoang et al. (2022). A full list of species names and accession number and sources can be found in Supplemental Table 1. Gas exchange data was obtained from Schlüter et al. (2022). The phylogenetic tree of all studied species was obtained from Guerreiro et al. (2023).

### Gene annotation

Consistent structural gene annotations were generated for each species using *Helixer* (Holst et al. 2023) with the hybrid convolutional and bidirectional long-short term memory model, `HybridModel`, specifically the trained instance of `land_plant_v0.3_m_0100` with default parameters.

### Annotation of transposable elements

*TEs were de novo* annotated using *EDTA* 1.9.9 (Ou et al. 2019) using the `-anno` 1 and `-sensitive` 1 flags. For the calculation of genomic composition (Fig. 2, Fig. 3), intact and fragmented TEs were used. To reduce the influence of false-positive hits, the pan-genomic gene-TE association study was performed for intact TEs only. The long terminal repeats (LTR) insertion time was calculated using

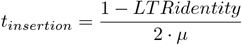

assuming a neutral mutation rate of *µ* = 1.4 *·*10^−8^ substitutions per site per year (Cai et al. 2018).

**Figure 2.**
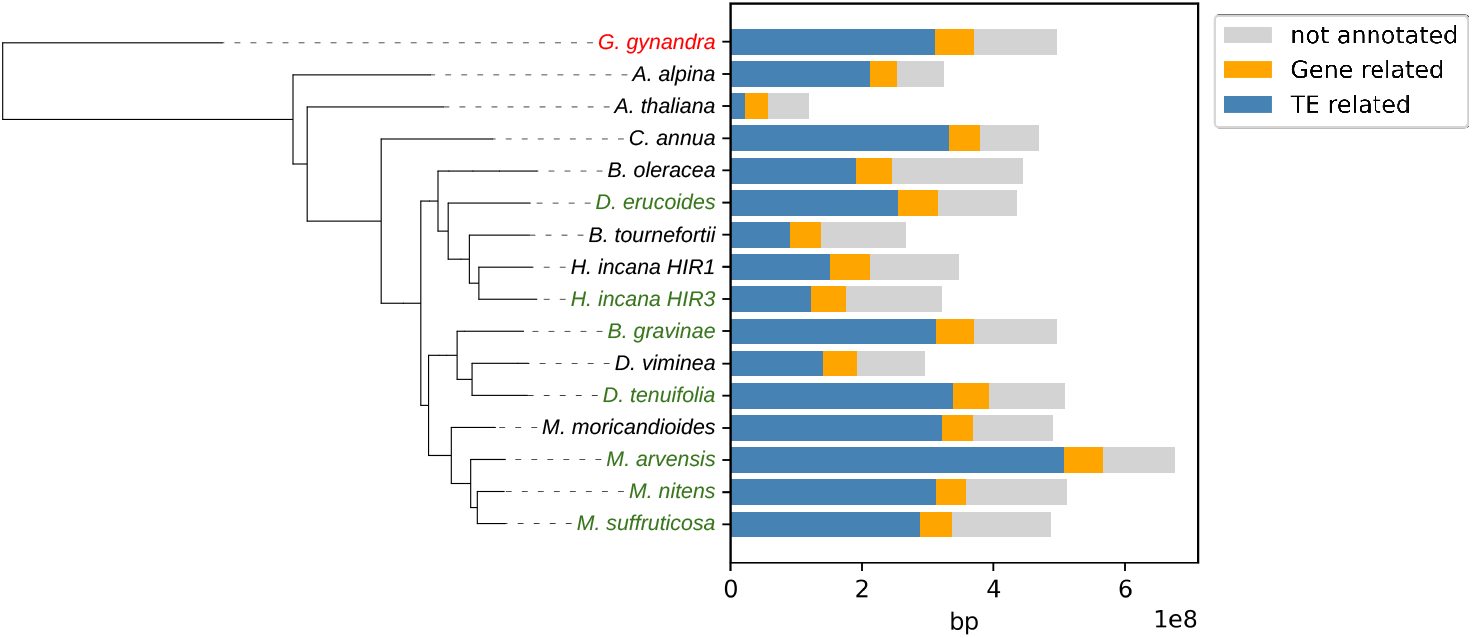
Phylogeny and genomic composition of 15 selected Brassicaceae species and the Cleomaceae outgroup. C_3_-C_4_ intermediate species are highlighted in green, the C_4_ outgroup *G. gynandra* is highlighted in red. Transposable element-related nucleotides are defined as spanning intact and fragmented transposons.

**Figure 3.**
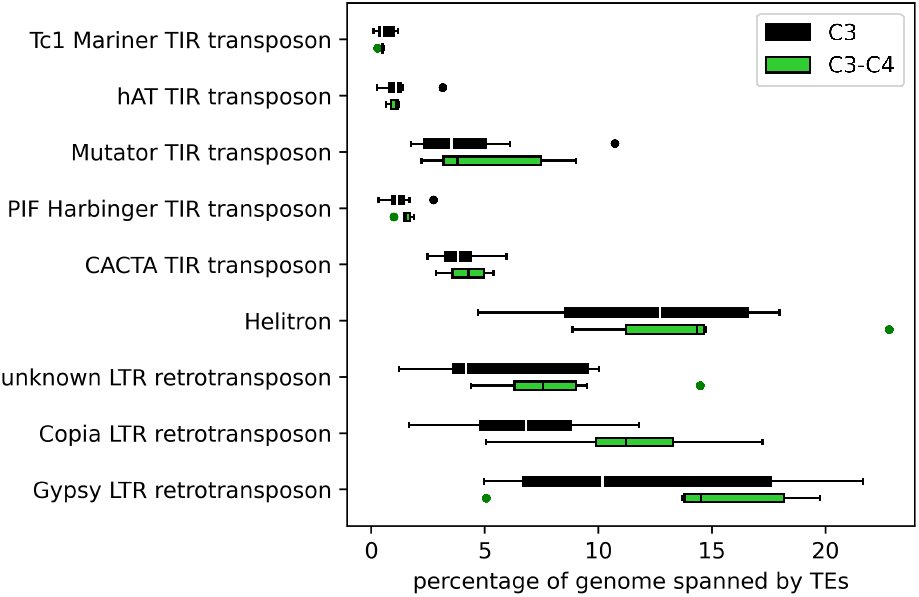
Boxplot indicating the percentage of the genome comprised by each class of intact and fragmented transposable elements (TEs) in eight C_3_ and six C_3_-C_4_ intermediate species. The y-axis shows the TE classes, the x-axis indicates the fraction of the genome made up by the respective TE class. Black boxes depict C_3_ species and green boxes depict C_3_-C_4_ intermediate species.

### Analysis of differential transposable element insertion

All downstream analyses were performed using *Python* 3.6 including *pandas* 1.2.4, *numpy* 1.20.1, *matplotlib* 3.4.1, *scikit-learn* 0.24.1, *scipy* 1.6.2 and *statsmodels* 0.12.2. All raw data and analyses are available in an Annotated Research Context (ARC) format under https://git.nfdi4plants.org/setri100/triesch2023_brassicaceae_transposons. The annotation files for genes and intact TEs were compared for each species. TEs were considered co-occurring with genes if their position matched one of the five cases described in Fig. 5. *CoGe SynMap* (https://genomevolution.org/coge/SynMap.pl) was used to identify orthologs and paralogs between the set of species. Each syntenic gene model was functionally annotated using *Mercator* 4.0 (Schwacke et al. 2019).

For each obtained syntelog, the effect of the presence or absence of an upstream TE on CCP was assessed using a phylogenetic implementation of the one-way ANOVA which accounts for the non-independence between species on the phylogenetic tree. For this purpose, phylogenetic ANOVAs were performed in the R environment using the *phylANOVA* function in the *phytools* 1.0.3 package (Revell 2012) using 1000 simulations and integrated posthoc comparisons to evaluate differences between means.

Enrichment of *Mercator* bins for genes with correlating upstream TEs was calculated using Fisher’s exact test. The identities of TEs in the *GLDP1* promoter were validated using the *CENSOR* webtool (Kohany et al. 2006).

## Results

To screen for genomic features of potential relevance to the evolution of the C_3_-C_4_ photosynthesis trait, we conducted a pan-genomic association study of eight C_3_ Brassicaceae species, seven C_3_-C_4_ intermediate Brassicaceae species from five independent origins, and one C_4_ Cleomaceae as an outgroup species for tree building. The five independent origins of C_3_-C_4_ intermediate photosynthesis can be found in the *Moricandia arvensis*, -*nitens*, and -*suffruticosa* monophylum, as well as in *Diplotaxis erucoides, Diplotaxis tenuifolia, Brassica gravinae* and *Hirschfeldia incana* HIR3 (Fig. 2) (Guerreiro et al. 2023, Schlüter & Weber 2016, Schlüter et al. 2022).

The species panel exhibits genome sizes ranging from 120 Mbp in *Arabidopsis thaliana* to 677 Mbp in *Moricandia arvensis*. We found no significant difference in genome size between species exhibiting either the C_3_ or C_3_-C_4_ intermediate photosynthesis phenotype (Fig 2; one-way ANOVA p*>*0.05). We next *de novo* annotated TEs using the *EDTA* pipeline (Ou et al. 2019). Overall, the annotated fragmented and intact transposons made up between 18 % of the genome in *Arabidopsis thaliana* and 75 % in *Moricandia arvensis*. We observed differences in genome size and TE content also in closely related species, between *M. arvensis* and *M. moricandioides* and between *B. gravinae* and *D. viminea*. Furthermore, we observed that differences in genome size are mainly due to the different TE content.

Class I type retrotransposons represented the majority of identified TEs across both C_3_ and C_3_-C_4_ species (Fig. 3). For instance, across all analyzed genomes, between 60 % and 68 % of all annotated TEs were Class I retrotransposons. In contrast, the proportion of TE classes in the genomes varied greatly across species (Fig. 3; Supplemental Table S2). The TE Class II was dominated by TEs from the *Helitron* group, making up between five and 20 percent of the genome (Fig. 3). The percentage of the genome made up of TEs from the different classes varied between the photosynthesis types, with a significantly higher amount of TEs in C_3_-C_4_ genomes (two-way ANOVA, p = 0.013).

To analyze recent increases of TE activity and their potential roles in the evolution of C_3_-C_4_ intermediate photosynthesis, we determined the insertion times of long terminal repeat (LTR) transposons (Fig. 4; Supplemental Table S3). *LTR retriever*, which is the LTR annotation tool of the *EDTA* pipeline, detected LTR transposons to a threshold for repeat identity of 91 %. Assuming a neutral mutation rate of µ =1.4*10^−8^ substitutions per site per year (Cai et al. 2018), LTR insertion times could thus be dated to a maximum of 4 million years ago. In general, both C_3_ and C_3_-C_4_ intermediate species revealed the same broad pattern of LTR bursts. Specifically, in both groups, there was an increased frequency for LTR-TEs younger than two million years. However, the increase was more pronounced for C_3_-C_4_ intermediate species, largely on account of the high number of young LTR-TEs in *M. arvensis*. Statistical analysis revealed a significant correlation between the age distribution of LTR-transposons and the photosynthesis phenotype (2-way ANOVA, p = 0.033).

**Figure 4.**
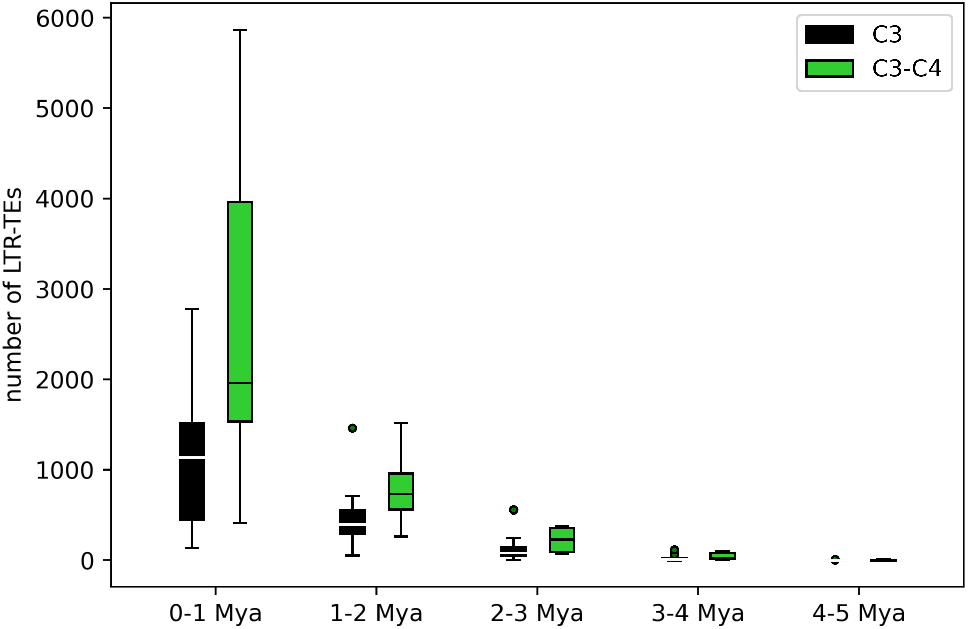
Boxplot of long terminal repeat transposable element (LTR-TE) insertion times for eight C_3_ and six C_3_-C_4_ intermediate species. The x-axis shows the insertion time in bins of 1 million years before today (Mya). The y-axis depicts the number of identified LTR-TEs calculated to be inserted within this timeframe. Calculation was performed using the LTR similarity of each LTR-TE and a neutral mutation rate of 1.4*10^−8^ substitutions per site per year. Black boxes represent C_3_ species, green boxes represent C_3_-C_4_ species.

To better understand whether the high abundance of TEs in C_3_-C_4_ species was global or associated with specific genes, we next analyzed the differential co-occurrence of TEs with protein coding genes. Co-occurrent TEs were defined as follows (Fig. 5): (I) the TE starts or ends in a 3,000 bp window upstream of the gene (upstream), (II) the TE starts or ends in a 3,000 bp window downstream of the gene (downstream), (III) the TE is residing within an exon or intron of the gene (inside), (IV) the TE starts but only partially resides in the gene (start), or (V) the TE ends but only partially resides in the gene (end).

**Figure 5.**
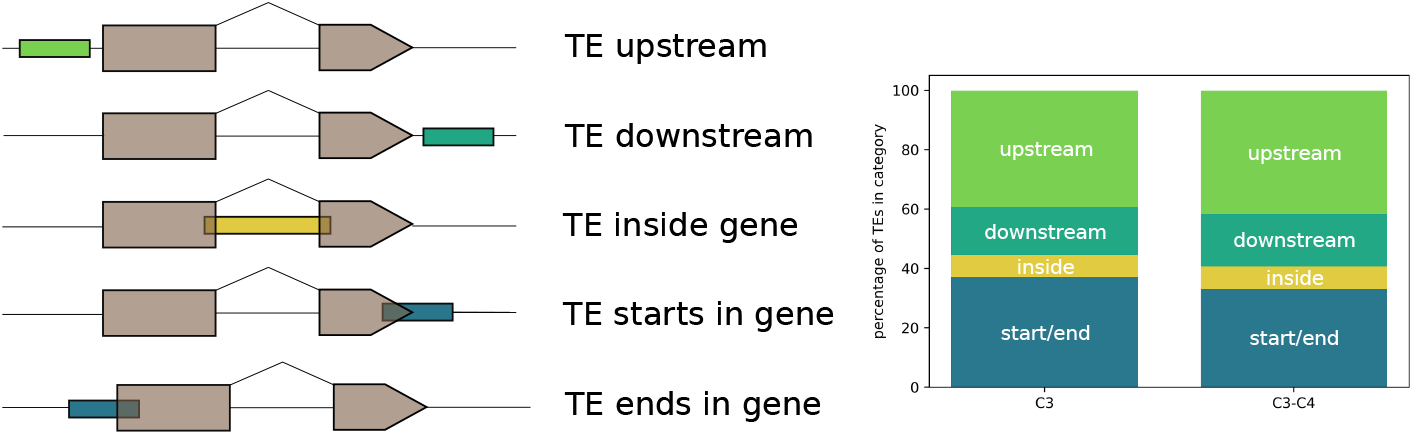
**Left panel:** Different contexts of transposable elements (TEs) co-occurring with genes. **Right panel:** Bar charts indicating the fractions of TE co-occurring with genes within five contexts: starting or ending in a gene (start / end), residing within a gene (inside) or residing within a 3000 bp window upstream or downstream the gene.

Genes with TEs within the gene model (III) overlapping TEs (IV and V) might have broken coding sequences and may result from imprecise annotations. Across the selected eleven species, 55,148 TEs were identified to be co-occurring with a protein coding gene in at least one species, whereas 21,643 co-occurring TEs belonged to C_3_ and 28,379 co-occurring TEs belonged to C_3_-C_4_ species. In both C_3_ and C_3_-C_4_ intermediate species, over 50 % of the TEs co-occurring with genes were located up- or downstream of the gene (Fig. 5). Analyzing potentially exaptated CREs, we focused on the up to 3000 bp 5 ‘ region of the gene. To compare differential TE insertions between the analyzed species, we obtained syntenic gene information for *CoGe SynMap*. For each of these syntenic gene models, one-way ANOVA was employed, correlating the presence or absence of a co-occurring upstream TE with the CCP of the respective species. After correcting the p-values for the phylogenetic bias, we identified 113 genes where the co-occurrence of one of the gene with an upstream TE correlated with the CCP (p *≤* 0.05; Tab. 1, Supplemental Table S4). Among the top ten genes (ranked by statistical confidence) were genes involved in photorespiration such as the genes encoding the T- and P-subprotein of the glycine decarboxylase complex (Fig. 6A). Strikingly, the C_3_-C_4_ intermediate orthologs of these genes exhibited upstream TEs, whereas the C_3_ orthologs lacked upstream TEs. Thus, during the evolution of C_3_-C_4_, there is a “gain “ in upstream TEs in genes that function in photorespiration (Fig. 6A). In the subset of genes which exhibit an association between the presence of an upstream TE and the plant CCP, two photorespiratory genes occurred (*GLDP, GLDT*). To quantify putative enrichment of certain gene ontologies, each gene was functionally annotated with a *Mercator* bin. Statistical enrichment analysis using Fisher’s exact test revealed that the *Mercator* bin “Photosynthesis.Photorespiration “ (p = 0.002907) was enriched in the set of genes that co-occur with upstream transposons (Tab. 2). The occurrence of this *Mercator* bin was increased 38-fold over the background, which is higher than for any other analyzed *Mercator* bin (Tab.2; Supplemental Table S5).

**Figure 6.**
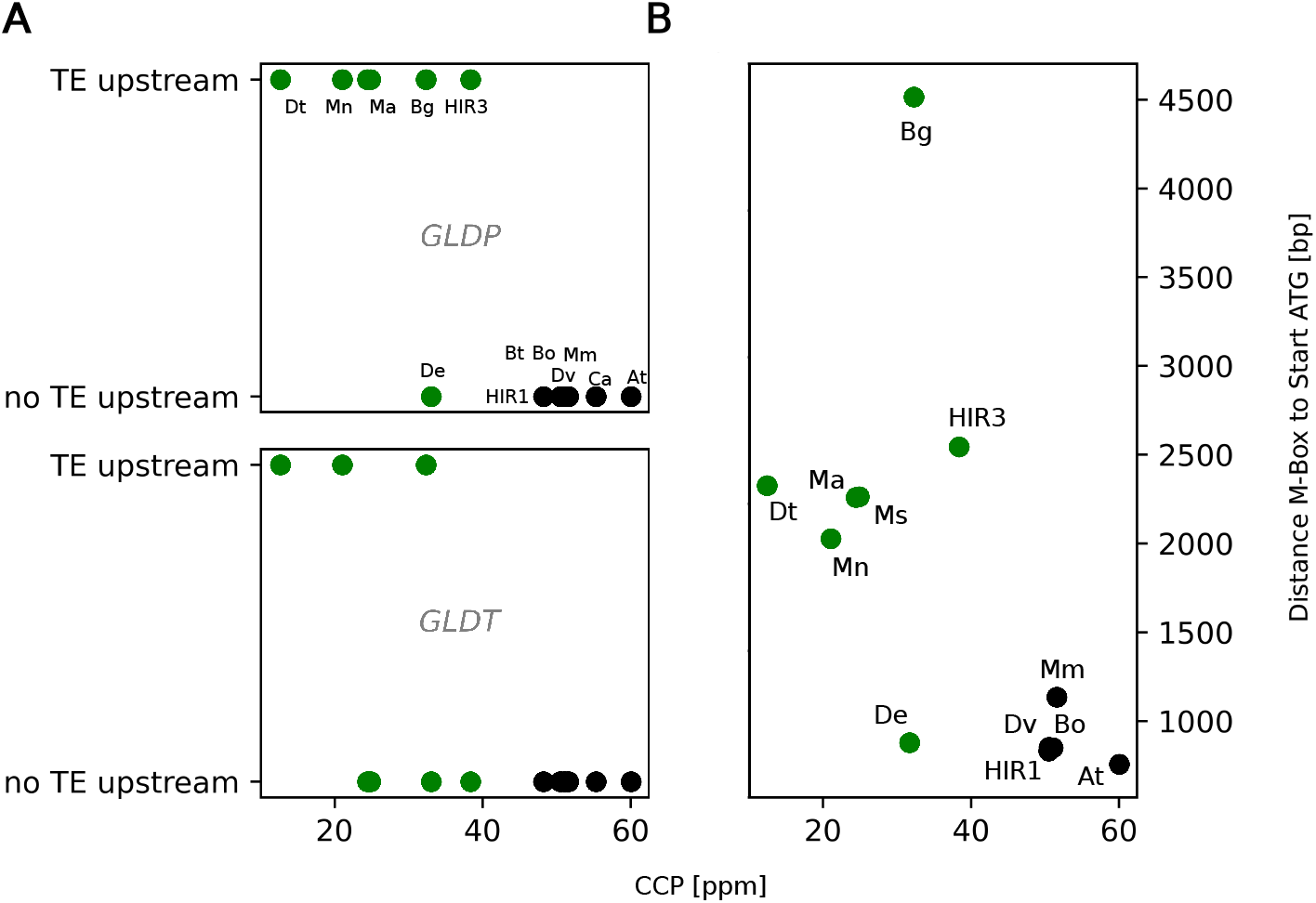
A: Scatter plot for two photorespiratory genes with significant co-associated upstream transposable elements (TEs.) The y-axis indicates the presence of an upstream TE (yes/no), the x-axis shows the carbon compensation point. Abbreviations: GLDP/GLDT: P/T-protein of the GLYCINE DECARBOXYLASE COMPLEX **B: Scatter plot for the different architectures of the *GLDP1* promoter**. The y-axis indicates the distance between the conserved M-Box sequence and the *GLDP1* start site. Each dot represents a species C_3_ species are shown in green, C_3_-C_4_ intermediate species are shown in black. Species name abbreviations: At.: *Arabidopsis thaliana*, Bg: *Brassica gravinae*, Bo: *Brassica oleracea*, Bt: *Brassica tournefortii*, Ca: *Carrichtera annua*, De: *Diplotaxis erucoides*, Dt: *Diplotaxis tenuifolia*, Dv: *Diplotaxis viminea*, HIR1: *Hirschfeldia incana* HIR1, HIR3: *Hirschfeldia incana* HIR3, Ma: *Moricandia arvensis*, Mm: *Moricandia moricandioides*, Mn: *Moricandia nitens*, Ms: *Moricandia suffruticosa*

As *GLDP* was the gene model with the strongest association between the presence of upstream TEs and CCP and it is known that the differential expression of *GLDP* contributes to the establishment of the photorespiratory glycine shuttle (Monson & Edwards 1984, Rawsthorne et al. 1988, Schulze et al. 2013), we selected this gene for further analysis. Several studies about the underlying regulatory genetics of *GLDP* expression have been conducted before (Adwy et al. 2015, 2019, Dickinson et al. 2020, Schulze et al. 2016). Only one *GLDP* gene copy is present in species from the Brassiceae tribe that contains all known C_3_-C_4_ intermediate species of the Brassicaceae (Schlüter et al. 2017). In contrast, the other two photorespiratory genes with correlating upstream TEs (Tab. 1; Fig. 6A) are found in higher copy numbers, which complicates a detailed genetic analysis.

We found three independent TE insertions in the promoter of C_3_-C_4_ intermediate *GLDP1* orthologs. In *Diplotaxis tenuifolia* a *Mutator* TE starts at 1970 bp upstream of the *GLDP1* start codon. In *Hirschfeldia incana* HIR3 a TE of the *Helitron* class is located around 2240 bp upstream. In orthologs from the monophyletic clade *Moricandia arvensis, Moricandia nitens* and *Moricandia suffruticosa* a *MITE* DNA transposon was detected, starting 1950 bp upstream of the *GLDP1* start codon. We calculated the minimum timespan since the *MITE* insertion by pairwise multiple sequence alignments of the *MITE* in the three *Moricandia GLDP1* promoters using the neutral mutation rate formula that was also employed for the calculation of LTR ages. We found the *GLDP1* promoter *MITE* was at least 6.5 million years old.

All three independent TE insertions are located around 100 bp downstream of the M-Box promoter motif. This motif was previously hypothesized to confer MC expression (Adwy et al. 2015) since truncation of the motif from the *AtGLDP1* promoter shifted GUS activity from the whole leaf apex to the veins. Furthermore, the M-Box was reported to be lost in C_3_-C_4_ intermediate *Moricandia* species (Adwy et al. 2019). However, upon closer inspection, we found a highly conserved M-Box motif in all Brassicaceae genomes analyzed here. Notably, the M-Box was shifted upstream due to the TE insertion in C_3_-C_4_ species with the exception of *D. erucoides* (Fig. 6B, Fig. 7, Supplemental Table S6). In *Brassica gravinae*, the *EDTA* pipeline did not annotate an upstream transposon. However, we found a large insertion of unknown origin in the *B. gravinae GLDP1* promoter. This insertion is larger than the three reported TE cases but could be found in a similar position compared to the other *GLDP1* promoter insertions of TE origin (Fig. 7). In the *GLDP1* promoter of C_3_-C_4_ intermediate species *D. erucoides* no insertion could be found.

**Figure 7.**
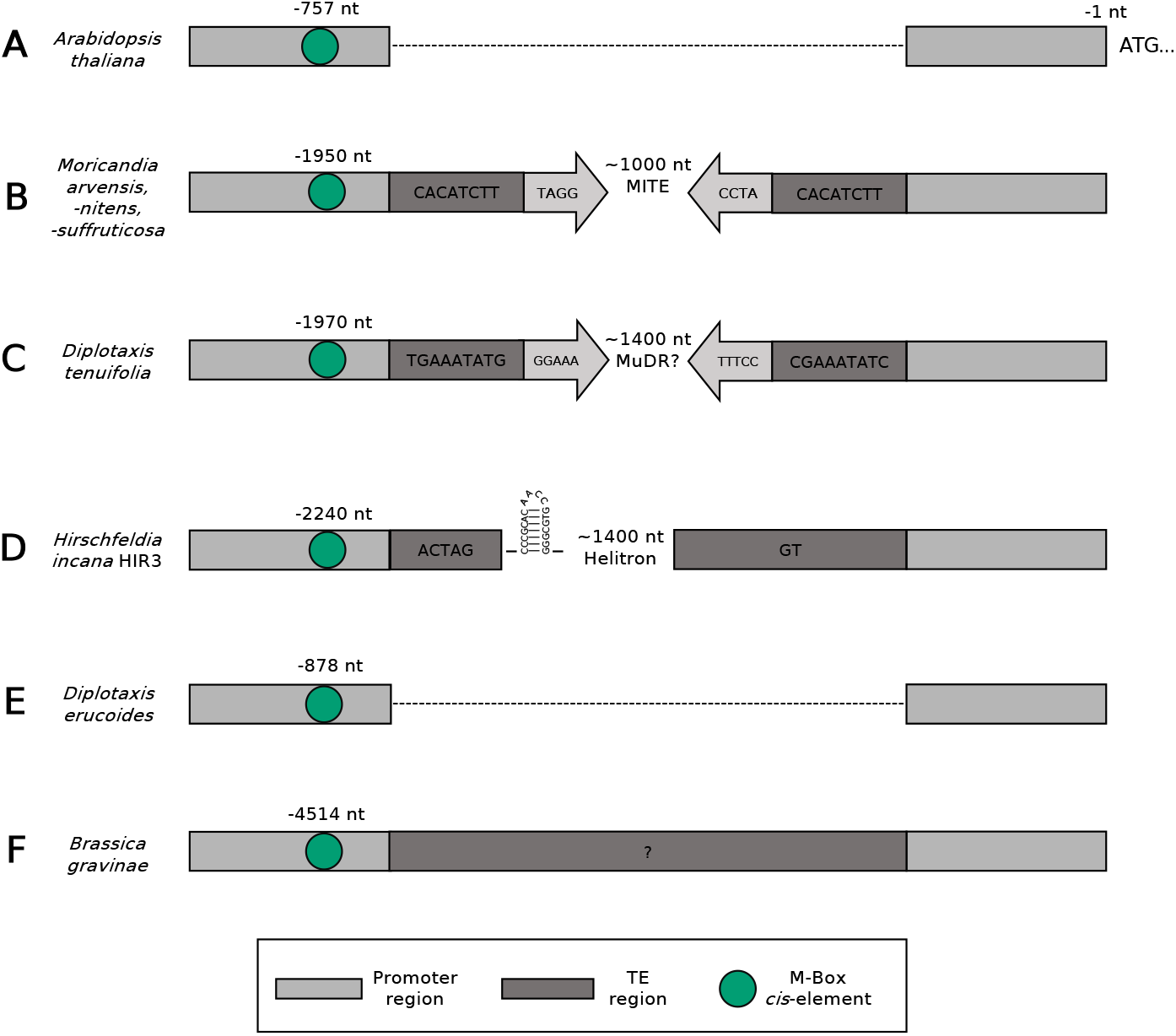
Schematic representation of the *GLDP1* promoter region. “ATG… “ depicts the start site of the *GLDP1* gene. Dark grey boxes represent characteristic transposable element (TE) sites such as target site duplications or the *Helitron* insertion sites. Grey arrows depict terminal inverted repeat motifs. The M-Box motif is highlighted as a green circle. In C_3_ species such as *Arabidopsis thaliana* no TE is annotated in the promoter sequence, leading to a low spacing between the M-Box and the *GLDP1* start site (**A**). In the C_3_-C_4_ intermediate *Moricandia* species, a *MITE* TE begins around 1950 bp upstream of the *GLDP1* start codon (**B**). In *Diplotaxis tenuifolia*, a *Mutator* TE begins 1970 bp upstream (**C**). In *Hirschfeldia incana* HIR3 a *Helitron* with a highly conserved hairpin loop structure is inserted around 2240 bp upstream (**D**). Although being a C_3_-C_4_ intermediate species, the *Diplotaxis erucoides GLDP1* promoter did not have an insertion behind the M-Box. (**E**). In *Brassica gravinae* a large insertion of unknown origin could be found behind the M-Box region (**F**).

From five analyzed C_3_-C_4_ *GLDP1* promoters we found a large insertion behind the conserved M-Box in four cases (monophyletic C_3_-C_4_ intermediate *Moricandia* clade, *Diplotaxis tenuifolia, Brassica gravinae* and *Hirschfeldia incana* HIR3; Fig. 6B). Out of these four cases where the insertions occurred, we found evidence for the sequence being a TE in three cases (Fig. 7).

## Discussion

Evolution of new complex traits such as C_3_-C_4_ photosynthesis and C_4_ photosynthesis requires the differential regulation of multiple genes. This includes differential gene expression across both MSC and BSC tissue as well as the installation of light-responsiveness for genes of the core metabolism (reviewed in Hibberd & Covshoff (2010)). In many cases, the evolution of differential gene regulation takes place in promoter sequences, either by introduction or suppression of *cis*-elements.

A few *cis*-elements for MC specificity have been previously found, including the MEM1 motif from the *Flaveria trinervia* phosphoenolpyruvate carboxylase gene (Gowik et al. 2017) as well as the M-Box sequence in Brassicaceae (Adwy et al. 2015, Dickinson et al. 2020).

TEs have the potential to deliver or suppress *cis*-elements upon their insertion in a target promoter. TEs can generate antisense transcription, interrupt or generate heterochromatic regions, or serve as raw material for the *de novo* evolution of new *cis*-elements (reviewed in Feschotte (2008)). The role of TEs in the evolution of C_4_ photosynthesis is only just started to being uncovered. The present study comprises the first pan-genomic association analysis to assess the importance of TEs in the evolution of C_3_-C_4_ intermediacy. Specifically, to do this, we analyzed the role of differential TE landscapes in 15 Brassicaceae species. Firstly, we investigated whether genome size and TE content correlate with the presence of the C_3_-C_4_ photosynthesis phenotype. Across our species panel a variety of genome sizes is present (Fig. 2), but we could detect no correlation between genome size and the presence of the photosynthesis trait. However, it is possible that different levels of heterozygosity in the sequenced species may confound these results and genome size estimations have to be handled with care.

Within the Brassicaceae family species exhibiting C_3_-C_4_ intermediate traits can only be found in the Brassiceae tribe. Notably, species from this tribe seem to have undergone recent polyploidization events (Walden et al. 2020) and exhibit larger genome sizes than species from neighboring tribes (Lysak et al. 2009).

Next, we analyzed the proportion of TEs across individual genomes. Our estimation of TE proportions is consistent with previously analyzed Brassicaceae genomes (Liu et al. 2020, Mirouze & Vitte 2014) and the *Gynandropsis gynandra* genome (Hoang et al. 2022). While genome size and TE content vary between species, we found a significant correlation between the photosynthesis phenotype and the proportion of the genome occupied by TEs in the respective species. Moreover, we found a recent burst in LTR-TE activity that is consistent with other studies (e.g. Cai et al. (2018)). The recent sharp increase in LTR-TE bursts in C_3_-C_4_ species comes mainly from *Moricandia arvensis* and might rather be due to high heterozygosity of LTR-containing genomic regions (Fig. 4). Although we found a significant correlation between LTR content and age with the C_3_-C_4_ intermediate phenotype, we cannot ultimately conclude that LTR transposon bursts contributed to the evolution of the C_3_-C_4_ intermediacy. Our LTR age analysis is limited to an LTR age of 4 million years. Given the estimated divergence time of 2-11 million years for C_3_ and C_3_-C_4_ intermediate *Moricandia* species (Arias et al. 2014), our analysis of LTR insertion times will miss the contribution of older LTRs to the evolution of C_3_-C_4_ intermediate traits. Furthermore, based on sequence identity between the C_3_-C_4_ intermediate *Moricandia GLDP1* promoters, we estimate the age of the *MITE* in the *Moricandia GLDP1* promoter to be at least 6.5 million years. This also falls within the proposed divergence time C_3_ and C_3_-C_4_ intermediate *Moricandia* species of 2-11 million years (Arias, 2014). Thus, changes in TE content occurred concomitant with the evolution of C_3_-C_4_ intermediate photosynthesis and occurred in genes whose expression is required to change for operation of a C_3_-C_4_ cycle.

In the descriptive whole-genome view, we observed correlations between TE content and age and the C_3_-C_4_ intermediate phenotype. Yet, however, there is an individual TE pattern even in closely related lines (Fig. 2). We therefore conclude that the role of TE activity may have an influence on C_3_-C_4_ evolution, but not necessarily via means of general TE activity (TE outbursts or TE purging) but rather via selective TE insertions to relevant genes or upstream regions. To analyze this, we employed a pan-genomic *de novo* transposon-gene association study, where we correlated the co-occurrence of TEs with genes to the presence of a C_3_-C_4_ intermediate phenotype.

In both C_3_ and C_3_-C_4_ intermediate species, more than 50 % of the analyzed co-occurring TEs were upstream or downstream of the respective co-occurring gene or spanning the gene. This is biologically plausible, as TEs crossing gene borders may disturb gene function and intergenic regions can harbor transposable elements (Buchmann et al. 2012). Nevertheless, we found over 30 % of the transposons crossing the borders of annotated genes. We assume that this was due to imprecise annotations by the TE identification pipeline. Differential gene regulation mediated by variation in upstream regions was shown to be a driver of C_4_ trait evolution in multiple, well documented cases (Adwy et al. 2015, Gowik et al. 2017, Williams et al. 2015, Wiludda et al. 2012). Our analysis revealed 113 genes with an upstream TE that correlates with the presence of a C_3_-C_4_ intermediate phenotype (Fig. 7; p*<*0.05). Enrichment analysis of *Mercator* bins for this set of genes revealed an enrichment of the codes “Multi-process regulation.sucrose non-fermenting-related kinase (SnRK1) regulation “ and “Photosynthesis.Photorespiration “. SnRK1 was shown to act as a central regulator of starvation metabolism that mediates energy homeostasis between organelles (Wurzinger et al. 2018). During nutrient starvation, SnRK1 subcomplexes were found to regulate the differential expression of over 600 target genes (Baena-González et al. 2007). Strikingly, ultrastructural adjustments and re-localization of the GDC P-protein to the BSC were demonstrated as a result of nitrogen starvation in the C_3_-C_4_ intermediate species Chenopodium album (Oono et al. 2022).

There is a clear bias of TE retention upstream of photorespiratory and SnRK1-regulatory genes in C_3_-C_4_ intermediate species, although with a small effect size (2 out of 5 genes with p*<*0.05 for “Photosynthesis. Photorespiration “; 2 out of 11 genes with p*<*0.05 for “Multi-process regulation.SnRK1 regulation “; see Table 2). We suggest that TE retention upstream of these genes has functional consequences such as differential gene expression, putatively due to the co-option of new, or, suppression of existing *cis*-elements. Strikingly, the set of genes that are significantly enriched for the presence of TEs in the upstream region contains multiple genes involved in photorespiration, such as those encoding the T- and P- proteins of the glycine decarboxylase complex (GLDT/GLDP). The modification of photorespiration is an important step towards the establishment of the glycine shuttle. The enrichment of TE insertions upstream of photorespiratory genes in C_3_-C_4_ intermediates is a potential hint that TEs play a significant role in the introduction of the glycine shuttle.

**Table 1.** Selected subset of ten genes with upstream transposable elements (TEs) with the lowest p-values for their association with the CCP.

| Gene Name | AGI locus code | p-value |
| --- | --- | --- |
| glycine dehydrogenase component P-protein of glycine cleavage system | AT4G33010 | 0.001 |
| negative on TATA-less (NOT2) | AT5G59710 | 0.003 |
| regulatory protein FLZ of SnRK1 complex | AT5G49120 | 0.004 |
| pectate lyase | AT5G63180 | 0.005 |
| MATE efflux family protein | AT2G38510 | 0.005 |
| CYCLIN D-type regulatory protein | AT4G34160 | 0.005 |
| regulatory protein FLZ of SnRK1 complex | AT5G47060 | 0.005 |
| phosphocholine phosphatase (PS2/PECP1) | AT1G17710 | 0.007 |
| PLATZ transcription factor family protein | AT3G50808 | 0.007 |
| U-box domain-containing E3 ubiquitin ligase | AT4G25160 | 0.007 |

**Table 2.** Results from two-sided Fisher’s exact test for the enrichment of *Mercator* bins within the set of genes with significant upstream transposons.

| *Mercator* Bin | genes with<br>p>0.05 | genes with<br>p<0.05 | p-value | Odds ratio |
| --- | --- | --- | --- | --- |
| Photosynthesis.Photorespiration | 3 | 2 | 0.002907 | 38.2 |
| Multi-process regulation.SnRK1-kinase regulation | 9 | 2 | 0.014932 | 12.7 |
| Cell wall organisation.cell wall proteins | 32 | 3 | 0.022505 | 5.4 |
| Solute transport.channels | 45 | 3 | 0.050587 | 3.8 |

*GLDP* is a well-characterized example for differential gene expression at the early stages of C_3_-C_4_ evolution across multiple plant lineages (Schlüter & Weber 2016, Schulze et al. 2013). In the Brassiceae tribe, the *GLDP2* copy was lost (Schlüter et al. 2017). Additionally, *GLDP1* was reported to be differentially expressed between C_3_ and C_3_-C_4_ intermediate *Moricandia* species (Hylton et al. 1988). In *A. thaliana*, GUS activity was restricted to the BSC by truncating the *GLDP1* promoter in the position of the M-Box, a promoter element approx. 800 bp upstream of the *AtGLDP1* gene start site. It was hypothesized that the M-Box confers MC expression, whereas the V-Box, a second promoter element shortly before the At*GLDP1* start site, confers BSC expression (Adwy et al. 2015). Promoter-*GUS* fusions showed that the *GLDP1* promoter of the C_3_ species *M. moricandioides* conferred *GUS* expression to both MC and BSC, whereas the *GLDP1* promoter of the C_3_-C_4_ intermediate species *M. arvensis* restricted *GUS* expression to the BSC (Adwy et al. 2019).

Adwy et al. (2019) explain the establishment of the glycine shuttle in *Moricandia* by the loss of the M-Box in C_3_-C_4_ intermediate *Moricandia* species. However, in contrast to this, we found the M-Box sequence in all our analyzed *GLDP1* promoter variants, though this motif was shifted by over 1000 bp further upstream by the insertion of three independent TEs in the promoters in three independent evolutionary origins of C_3_-C_4_ intermediate photosynthesis, and by an insertion of unknown provenance in a fourth independent origin. This shift may have led to the M-Box being overlooked in previous studies.

Based on the findings by Adwy et al. (2019) we conclude that not the loss of the M-Box, but rather the upstream shift of the element by insertion of a TE has led to the differential tissue specific expression of the *GLDP1* gene. The upstream shift of the M-Box was mediated by three independent TE insertions in lines with independent evolutionary origins of C_3_-C_4_ photosynthesis. This hints at a remarkable convergent evolutionary genetic mechanism in C_3_-C_4_ evolution. We suggest that the loss of *GLDP2* paved the way for neofunctionalization of the *GLDP1* copy in the Brassiceae tribe, the only Brassicaceae tribe containing C_3_-C_4_ intermediate species. This was mediated by the insertion of a TE in the promoter, suppressing the M-Box element and shifting *GLDP1* expression. It is questionable whether the TE insertion took place before or after the preconditioning of C_3_-C_4_ photosynthesis by anatomical adaptations such as higher vein density and the distinct leaf anatomy. Hypothetically, limited expression of *GLDP1* in the MC may have been deleterious without further adaptations, which could have prevented the TE retention in the promoter. In *D. erucoides* we do not find a transposon in the *GLDP1* promoter region. The spacing of the M-Box to the *GLDP1* start codon is in the range of C_3_ plants (Fig. 6B). However, *D. erucoides* shows C_3_-C_4_ intermediate phenotypes (Lundgren 2020, Schlüter et al. 2017). We assume that, being an independent evolutionary origin of C_3_-C_4_ intermediate photosynthesis, *D. erucoides* either shifted *GLDP1* expression to the BSC by different means or, alternatively, that there must be other additional regulators in the *GLDP1* promoter beyond our transposon-M-Box model. Contrasting the well-studied GDC activity and localization in *Moricandia* species, there is no data on the *D. erucoides* GDC biochemistry and genetics. Therefore, we cannot rule out that the glycine shuttle in *D. erucoides* is mediated by a different GDC regulation compared to the other C_3_-C_4_ intermediate species, such as the differential activity of the GDC T-, L- or H-proteins.

By adopting a whole-genome view of TE density and gene-TE associations, our study highlights the potential importance of TE insertions in contributing to the convergent evolution of C_3_-C_4_ intermediacy. Differential *GLDP* expression is one of the most important innovations that occurs and facilitates the establishment of the glycine shuttle. The novel genetic mechanism of differential *GLDP1* regulation by a TE-mediated insertion causing an upstream shift of the M-Box must be verified in experimental work. Looking at the genetic mechanisms of gene regulation in C_3_-C_4_ intermediate species will pave the way for a better understanding of the C_4_ trait and facilitate genetic engineering efforts.

## Acknowledgements

This work was funded by the Deutsche Forschungsgemeinschaft (German Research Foundation) under Germany ‘s Excellence Strategy EXC-2048/1 under project ID 390686111, the Deutsche Forschungsgemeinschaft under Project ID 391465903/GRK 2466, the ERA-CAPS (European Research Network for Coordinating Action in Plant Sciences) project C4BREED under Project ID WE 2231/20–1 and the CRC (Collaborative Research Center) TRR341 under Project ID 456082119. JWB was funded by the BBSRC through BB/J014427/1. SK was funded by a Royal Society University Research Fellowship.

## Author contributions

A.P.M.W., B.S. and U.S. designed and coordinated the project. S.T. designed and integrated all analyses. J.W.B and S.K. performed the phylogenetic correction of p-values. N.B. performed synteny analysis using *CoGe SynMap*. A.K.D. performed gene annotations using *Helixer*. A.K.D., R.N.F.M.G. and B.S. advised on statistical testing. All authors contributed to writing and accepted the manuscript.

